# Detection of known and novel virus sequences in the black solider fly and expression of host antiviral pathways

**DOI:** 10.1101/2024.03.29.587392

**Authors:** Hunter K. Walt, Heather R. Jordan, Florencia Meyer, Federico G. Hoffmann

## Abstract

Mass rearing of animals in close quarters can be highly conducive to microbe transmission, including pathogens. This has been shown multiple times in the case of important industrial insects such as crickets, silkworms, and honeybees. One industrial insect of increasing importance is the black soldier fly (Diptera: *Hermetia illucens*), as it can convert organic waste into high quality protein and fatty acids. Along with this, they take up far less space than traditional protein sources, as millions of black soldier flies can be reared in a relatively small facility. Because of this, there is a growing interest in the pathogens that could impact black soldier fly rearing efforts. So far, only three black soldier fly-associated viruses have been identified. We used metatranscriptomic sequencing to survey black soldier fly guts, frass, and diet for viruses. We detected sequences from two novel viruses. One, which we name *Hermetia illucens* sigma-like virus 1, is phylogenetically related to viruses of the genus Sigmavirus, which have been highly studied in *Drosophila*. The other novel virus, which we name *Hermetia illucens* toti-like virus 2, is the second toti-like virus to be described in the black soldier fly. We also detected two black soldier fly-associated viruses previously identified by our group: BSF nairo-like virus, and BSF uncharacterized bunya-like virus. Consistent with our previous study, these two viruses are found primarily in frass samples and occur together more often than expected at random. When analyzing host transcription, we found significant differences in gene expression for eight candidate antiviral genes in black soldier fly when comparing samples with and without viral sequences. Our results suggest that black soldier fly-virus interactions are ongoing, and they could be of interest to black soldier fly producers.

## 1. INTRODUCTION

Insects provide an attractive alternative to traditional protein sources as a more sustainable and environmentally friendly option. The black soldier fly (Diptera: *Hermetia illucens*) (BSF) is a promising candidate as an alternative source of nutrition for livestock, pets and humans (Tomberlin & van Huis, 2020; van Huis, 2020). This is mainly due to the ability of BSF to convert organic waste to high quality proteins and fatty acids. Another appeal of BSF as an alternative protein source is that millions of BSF can be reared in a relatively small amount of space. Mass rearing of animals creates an opportune environment for the spread of microbes throughout a population. In the case of pathogens, this can be detrimental to an industry, and the production other insects of industrial importance has been affected by viral epidemics in the past (Maciel-Vergara & Ros, 2017).

Until recently there has not been a manifested interest in the viruses harbored by BSF. This is partially because BSF are believed to have a highly robust immune system due to the fact that the BSF larval stage lives in compost, and their genome encodes some of the highest numbers of antimicrobial peptide (AMP) genes in insects (Moretta et al., 2020; Vogel et al., 2018; Zhan et al., 2020). So far, two studies have detected viruses in BSF using shotgun metatranscriptomic approaches, and they have described three viruses associated with BSF, one toti-like virus and two bunyaviruses (Pienaar et al., 2022; Walt et al., 2023). Although the fitness effects of these viruses on BSF has not been assessed, one study detected many endogenous viral elements (EVEs) in the BSF genome, suggesting that BSF has historically been infected by viruses (Pienaar et al., 2022).

Because the BSF virome is poorly characterized and the interest in BSF production is increasing worldwide, we used a shotgun metatranscriptomic approach to detect viruses associated with BSF. We surveyed 76 larval gut (n=36), frass (n=36), or diet (n=4) transcriptome samples and detected two known and two novel BSF-associated viruses. Interestingly, one of the novel viruses is related to a *Drosophila* virus that has been routinely studied as a host-pathogen coevolution model in *Drosophila*, and the other adds to a growing clade of insect-associated toti-like viruses. These viruses could be of interest to BSF producers, as they may affect BSF fitness. Finally, we found that multiple candidate antiviral genes were differentially expressed in samples where viral sequences were detected.

## 2. MATERIALS AND METHODS

### 2.1 BSF and Substrate Sampling

BSF larvae were sampled from an experiment determining the performance of BSF larvae reared on four different diet substrates of varying nutritional profiles. Briefly, larvae were reared in 50 ml conical plastic tubes and fed either Gainesville diet (industry standard), a protein-biased diet (1C:5P), carbohydrate-biased diet (5C:1P), or a balanced diet (1C:1P). After the feed trial, larvae and frass samples were stored in a freezer at -80°C. We sampled nine larvae from each of the four diets (n=36), and briefly submerged them in 70% ethanol followed by a sterile water rinse. We dissected whole larval guts in RNAlater solution (Invitrogen, Waltham, MA, USA) in a sterile petri dish using a SteREO Discovery microscope (Zeiss, Oberkochen, Germany) and immediately placed them in 1.5 mL of RNAzol (Molecular Research Center, Cincinnati, OH, USA) for RNA isolation. Concordantly, we sampled 1 gram of substrate (frass) from each enclosure and placed it in 1.5 mL of RNAzol for RNA isolation. Samples of diet without larvae were run alongside the feed trial study, so we also sampled 1 gram of diet from each diet-only enclosure (n=4) and placed it into 1.5 mL RNAzol for RNA isolation.

### 2.2 RNA Isolation

All samples were homogenized in RNAzol using a Genolyte 1200 tissue homogenizer (Spex SamplePrep, Metuchen, NJ, USA) at 4,000 RPM for one minute using sterile 3mm stainless steel balls. The resulting homogenate was spun down for five minutes at 12,000G, and 1 mL of the supernatant was used for RNA isolation. The manufacturer’s protocol for RNA isolation using RNAzol was followed through DNA, protein, and polysaccharide precipitation, upon which the aqueous fraction containing the RNA was directly placed into a NEB Monarch RNA cleanup kit (New England Biolabs, Ipswich, MA, USA), also following the manufacturer’s protocol. RNA purity was measured using a NanoDrop One spectrophotometer (Thermo Fisher Scientific, Grand Island, USA), and RNA concentration was assessed using a Qubit 2.0 fluorometer (Invitrogen, Waltham, MA, USA). RNA integrity was assessed using a 4150 TapeStation System (Agilent, Santa Clara, CA, USA).

### 2.3 Library Preparation and Sequencing

Whole shotgun metatranscriptome sequencing libraries were prepared using NEB Ultra II RNA kits, but rRNA depletion and mRNA enrichment steps were avoided to keep all host and microbial reads. Resulting cDNA was multiplexed using NEBNext Oligos for Illumina (New England Biolabs, Ipswich, MA, USA). The resulting libraries were pooled by diet and the quality of the library was assessed using a 4150 TapeStation System (Agilent, Santa Clara, CA, USA). The resulting libraries were sequenced on a HiSeq 2000 instrument (Illumina, San Diego, CA, USA), generating 151 base pair reads.

### 2.4 Metatranscriptome Assembly

The quality of the reads was assessed using fastQC v.0.11.9 (Andrews, 2010) and adapters and low-quality bases were trimmed using Trimmomatic v.0.39 (Bolger et al., 2014). Black soldier fly reads were discarded from further analyses by mapping all sequencing datasets to the black soldier fly reference genome (GCF_905115235.1) (Generalovic et al., 2021) using HISAT2 v.2.2.1 (Kim et al., 2019) and redirecting the unmapped reads to a new file. For each dataset, the unmapped reads were assembled using Trinity v.2.14.0 through the Trinity docker container (Grabherr et al., 2011; Merkel, 2014). All transcripts were uniquely named across every transcriptome and all transcriptomes were concatenated into one file. Only transcripts greater than 500 nucleotides were retained and these were clustered at a minimum sequence identity of 90% using cd-hit-est (Li & Godzik, 2006).

### 2.5 Identification of Virus Sequences

All clustered transcripts were aligned to NCBI’s non-redundant protein database using DIAMOND in the BLASTX mode employing the –very-sensitive option (Buchfink et al., 2015, 2021). All sequences with alignments to known RNA viruses were further inspected for the presence of open reading frames using NCBI’s orffinder tool (https://www.ncbi.nlm.nih.gov/orffinder/) and the resulting proteins were analyzed for conserved protein domains using NCBI’s conserved domain database search tool (https://www.ncbi.nlm.nih.gov/Structure/cdd/wrpsb.cgi) and the InterProScan web server (Quevillon et al., 2005).

### 2.6 Phylogenetic Analysis of Novel Virus Sequences

To assess the identity of potentially novel viruses, we conducted phylogenetic analyses based on viral RNA-dependent RNA polymerase (RdRp). We collected a diverse set of viral RdRp sequences based on predicted viral order from conserved protein domain searches, and BLASTX searches (https://blast.ncbi.nlm.nih.gov/Blast.cgi) against NCBI’s non-redundant protein database. We aligned the amino acid sequences using the g-ins-i algorithm in MAFFT v.7.490 (Katoh & Standley, 2013). To build the phylogenetic trees, we used IQ-TREE v. 2.0.7 using MODELFINDER to identify the best-fitting model (Kalyaanamoorthy et al., 2017; Minh et al., 2020; Nguyen et al., 2015). Branch support was evaluated using the ultrafast bootstrap algorithm (Hoang et al., 2018; Minh et al., 2013). Resulting phylogenetic trees were viewed and annotated using the Interactive Tree of Life web server (Letunic & Bork, 2021).

### 2.7 Overlap of Virus Occurrence Across Samples

To test if the amount of co-occurrence in viruses across samples was more than one would expect at random, we used Fisher’s Exact Test implemented in R (R Core Team, 2020).

### 2.8 Quantification of BSF and viral transcripts

To determine if viral infection influenced putative antiviral gene expression in BSF, we pseudoaligned the trimmed read datasets from larval samples to the RefSeq BSF transcriptome using Salmon v0.14.1, using the reference genome as a decoy (Patro et al., 2017). We imported the abundance estimates from Salmon to DESeq2 (Love et al., 2014) using tximport (Soneson et al., 2015). One sample was discarded due to a low number of reads, and we exported normalized read counts from DESeq2.

To quantify viral transcripts, we mapped the trimmed datasets to the longest transcripts from *Hermetia illucens* toti-like virus 2 and *Hermetia illucens* sigma-like virus 1 using Kallisto v0.44.0 (Bray et al., 2016). We did not map to BSF uncharacterized bunya-like virus or BSF nairo-like virus as these viruses were primarily detected in frass samples, thus non-contiguous reads present in larval samples could be the result of contamination instead of viral infection in the BSF gut. We considered viruses “present” in a sample when more than 10 reads mapped to either virus transcript.

### 2.9 Identifying candidate antiviral genes in BSF

To identify candidate antiviral genes, we conducted a literature search to find relevant antiviral genes in *Drosophila* and mosquitoes (Rosendo Machado et al., 2021). We used these sequences to find orthologous genes present in the BSF genome using reciprocal blast hits. Along with this, we used eggNOG mapper v.2.1.9 in DIAMOND mode to assign KEGG identifiers to BSF genes putatively belonging to the major antiviral/immune pathways Toll, Imd, JAK/STAT, RNAi, and piRNA. Finally, we used the sequences of BSF AMPs and lysozymes identified by Vogel et al. (2018) to BLAST against the BSF genome and obtained the gene names for all putative AMPs/lysozyme genes in BSF. Using these genes, we tested for significant differences in putative antiviral BSF gene expression using the normalized read counts from DEseq2, and the Wilcoxon rank-sum test implemented in R. Genes with mean normalized counts less than 10 across all samples were not considered.

## 3. RESULTS

After filtering and clustering of our metatranscriptome assemblies, the total number of reads went from 4,315,370 transcripts to 381,196. Using BLASTX, we detected sequences from two known BSF-associated viruses (Walt et al., 2023) and two novel virus sequences in our metatranscriptomic assemblies.

### 3.1 Novel Virus Sequences

We detected two novel virus sequences in our study. One sequence had multiple BLAST alignments (E value = 0) to proteins belonging to the sigmaviruses. From now on, we refer to this sequence as *Hermetia illucens* sigma-like virus 1. To investigate phylogenetic relationships of this virus to other sigmaviruses, we aligned the RdRp of *Hermetia illucens* sigma-like virus 1 to a selection of sigmavirus RDRP sequences from RefSeq along with its top five closest BLAST hits. We used Rabies virus as an outgroup based off a previous study showing that it branches sister to the sigmaviruses in the rhabdovirus phylogenetic tree (Longdon et al., 2010). Our phylogeny shows that *Hermetia illucens* sigma-like virus 1 groups within a clade of dipteran-infecting sigmaviruses, including a *Drosophila melanogaster* sigmavirus and multiple louse fly-infecting sigmaviruses (**Figure 1A**). The genome of *Hermetia illucens* sigma-like virus 1 was fractionated across two transcripts, one 3,116 nt transcript containing the sigmavirus nucleocapsid protein gene (N), the putative polymerase-associated protein gene (P), and a partial matrix protein (M), and another 9,544 nt transcript containing three other canonical sigmavirus genes: the complete M protein gene, the spike protein (G) gene, and the RDRP-containing L gene (**Figure 1B**).

**Figure 1:**
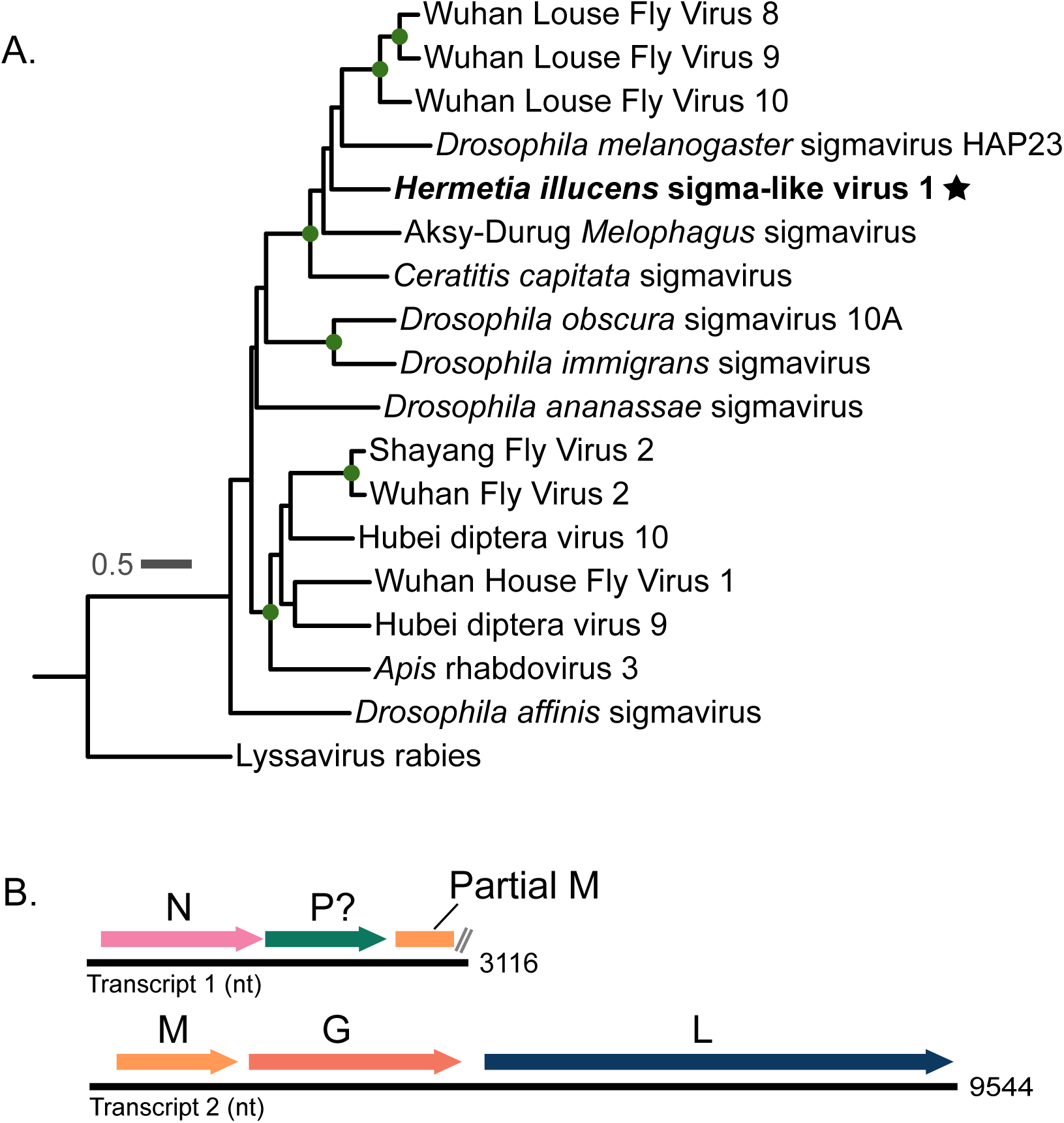
Sigmavirus phylogeny and genomic structure of *Hermetia illucens* sigma-like virus 1. **A.)** Phylogeny of the sigmaviruses, including *Hermetia illucens* sigma-like virus 1 which is denoted by a star. Green dots along the branches represent highly supported nodes: SH-alrt ≥ 80%, abayes ≥ 0.9, and ≥ 95% UF bootstrap. Another virus from the family Rhabdoviridae, lyssavirus rabies, was used as the outgroup **B.)** Genomic structure of *Hermetia illucens* sigma-like virus 1 according to our study. The question mark next to the P gene denotes that we could not confirm homology by sequence similarity. The fragmented genome could be a result of metatranscriptomic assembly error.

Along with *Hermetia illucens* sigma-like virus 1, we detected a transcript that had significant BLAST alignments to viruses of the family Totiviridae. We hereon refer to this sequence as *Hermetia illucens* toti-like virus 2. We aligned the translated RDRP region of this transcript with a large set of totivirus and toti-like virus RDRP amino acid sequences downloaded from NCBI (**Figure 2A**). We found that *Hermetia illucens* toti-like virus 2 groups in a highly supported clade of insect-infecting toti-like viruses, and branches sister to two toti-like viruses that were also detected in insects of the order Diptera (**Figure 2B**). The *Hermetia illucens* toti-like virus 2 genome is a 5,843 bp containing two large ORFs: one encoding a capsid protein, and one encoding the RDRP protein (**Figure 2C**).

**Figure 2:**
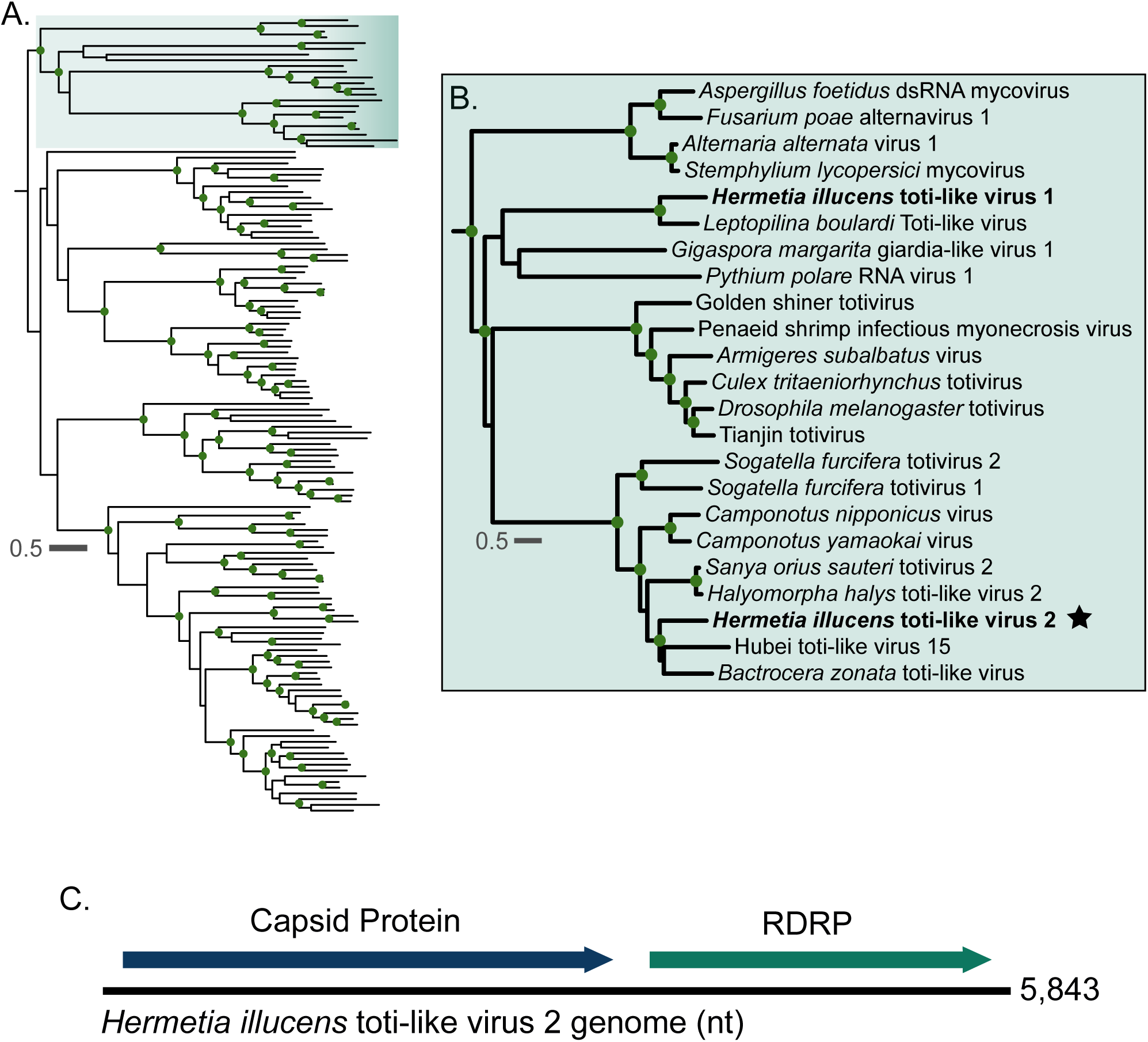
*Hermetia illucens* toti-like virus 2 groups within a highly supported clade of other dipteran toti-like viruses. **A.)** Full midpoint-rooted phylogeny of a diverse set of totiviruses and toti-like viruses. Green dots along the branches represent highly supported nodes: SH-alrt ≥ 80%, abayes ≥ 0.9, and ≥ 95% UF bootstrap. The green box denotes the area of the tree that is magnified in panel B. **B.)** Magnification of area of interest in panel **A.** BSF viruses are highlighted in bold, and BSF viruses discovered in this study are denoted by a star. **C.)** Genome structure of *Hermetia illucens* toti-like virus 2.

*Hermetia illucens* sigma-like virus 1 was detected in 11 samples from our study, all of which were larval gut samples (**Figure 3**). *Hermetia illucens* toti-like virus 2 was found in five total samples with four deriving from larval gut metatranscriptomes, and one deriving from a frass metatranscriptome (**Figure 3**).

**Figure 3:**
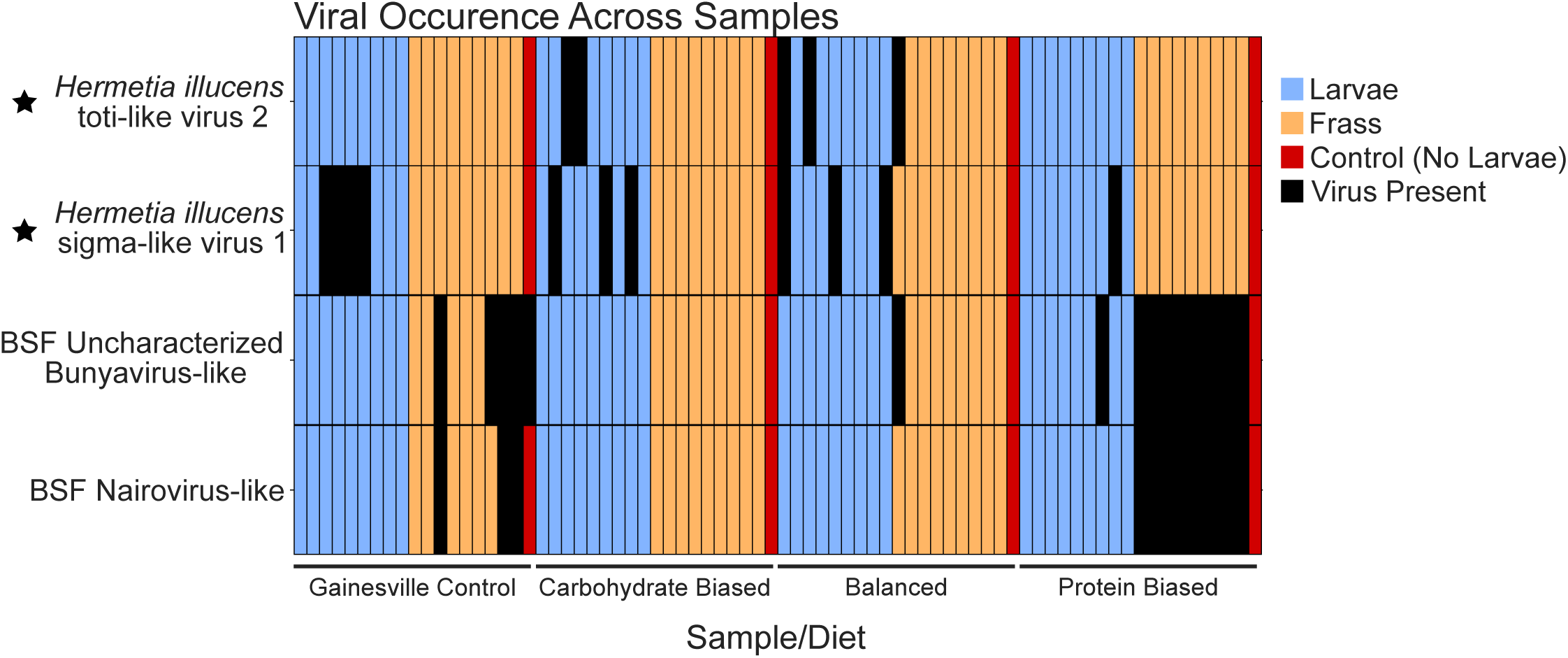
Occurrence of viruses across all samples used for this study. Novel viruses detected in this study are denoted by a star. BSF uncharacterized bunya-like virus and BSF nairo-like virus occur together more often than expected at random (Fisher’s Exact Test: p = 5.016e^-12^).

### 3.2 Known BSF-Associated Viruses

We detected two previously discovered virus sequences associated with BSF: BSF Uncharacterized bunya virus-like, and BSF nairo-like virus. These viruses primarily were detected in frass samples, similar to when we identified them (Walt et al., 2023) (**Figure 3**). BSF uncharacterized bunyavirus-like 1 occurred in 16 samples, with only one being derived from BSF larval gut (**Figure 3**). BSF nairo-like virus was detected in 12 samples, all of which were frass samples (**Figure 3**). Interestingly, these two viruses occur very regularly together. Every time BSF Nairo-like virus is detected, it co-occurs with BSF uncharacterized bunya-like virus, making them occur together 12 out of 16 times that one is present. This is significantly more than one would expect at random (Fisher’s Exact Test, p = 5.016e^-12^).

### 3.3 Antiviral gene expression in BSF

We identified 210 candidate antiviral genes from the dipteran immune/antiviral pathways Toll, Imd, JAK/STAT, RNAi, and piRNA using KEGG or literature search and BLAST (**Supplementary Table 1**). We tested all these genes for significant differences in expression between BSF samples that were classified as “virus-present” (greater than 10 reads mapped to a virus genome) relative to samples classifies as “virus absent” (less than 10 reads mapped to a virus genome). We found eight genes with significant differences in expression: three BSF orthologs associated with the Imd pathway in other dipteran insects, three associated with the JAK/STAT pathway in other dipterans, and two BSF AMPs (**Figure 4**). These genes along with the function of their *Drosophila* orthologs are shown in **Table 1**.

**Figure 4:**
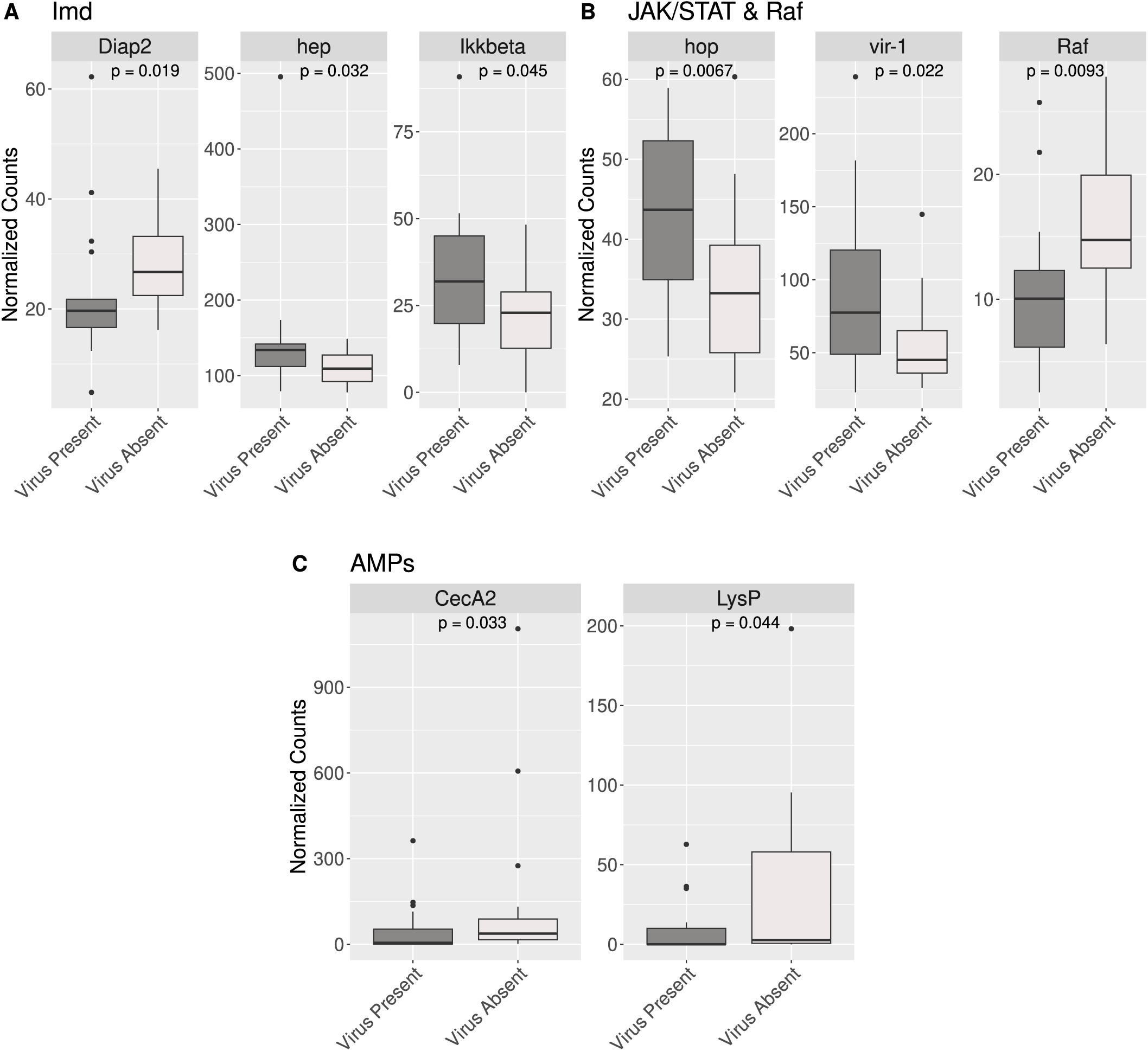
Eight candidate antiviral genes have significant differences in gene expression between virus-present BSF samples and virus-absent BSF samples. **A.)** Three genes putatively from the Imd pathway in BSF had significant differences in gene expression in virus-present vs. virus-absent BSF. Diap2 was upregulated in virus-absent samples, while hep and IKKbeta were upregulated in virus-present BSF samples. **B.)** Two genes from the JAK/STAT pathway were upregulated in virus-present BSF samples. Along with these, Raf, a gene whose product has known interactions with the JAK/STAT pathway was upregulated in virus-absent BSF samples. **C.)** Two AMPs were significantly upregulated in virus-absent samples. All gene names are derived from their ortholog in *Drosophila melanogaster*.

**Table 1:**
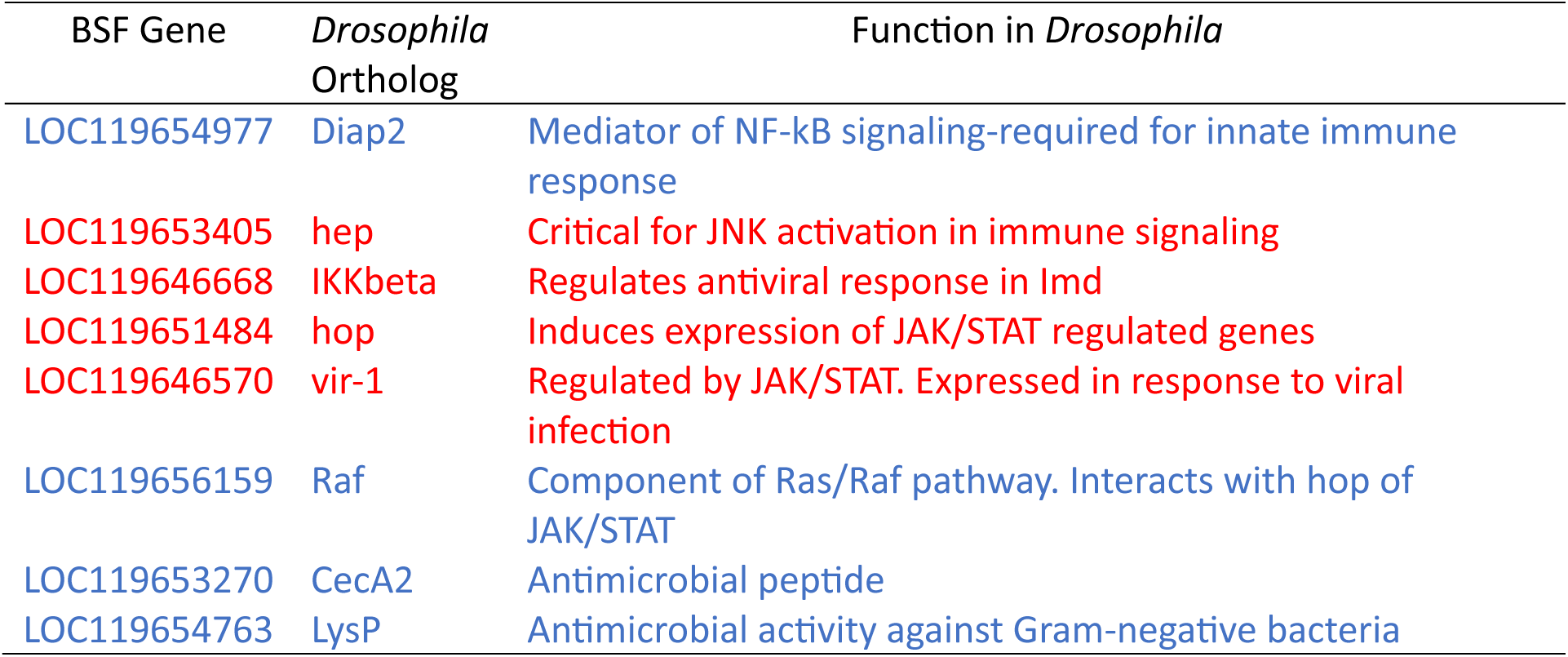
Significantly differentially expressed BSF genes and the function of their *Drosophila melanogaster* orthologs. Genes that are downregulated in virus-present samples are shown in blue text, while genes that are upregulated in virus-present samples are in red text.

| BSF Gene | *Drosophila*<br>Ortholog | Function in *Drosophila* |
| --- | --- | --- |
| LOC119654977 | Diap2 | Mediator of NF-kB signaling-required for innate immune response |
| LOC119653405 | hep | Critical for JNK activation in immune signaling |
| LOC119646668 | IKKbeta | Regulates antiviral response in Imd |
| LOC119651484 | hop | Induces expression of JAK/STAT regulated genes |
| LOC119646570 | vir-1 | Regulated by JAK/STAT. Expressed in response to viral infection |
| LOC119656159 | Raf | Component of Ras/Raf pathway. Interacts with hop of JAK/STAT |
| LOC119653270 | CecA2 | Antimicrobial peptide |
| LOC119654763 | LysP | Antimicrobial activity against Gram-negative bacteria |

## 4. DISCUSSION

### 4.1 *Hermetia illucens* sigma-like virus 1

Our study identifies two novel BSF-associated viruses and two previously described BSF-associated viruses. Before this, only three exogenous viruses had been found in BSF (Pienaar et al., 2022; Walt et al., 2023). One of the viruses discovered in this study, which we call *Hermetia illucens* sigma-like virus 1, is phylogenetically related to the sigmaviruses (**Figure 1**). Sigmaviruses are a genus of negative-sense single-stranded RNA viruses within the family Rhabdoviridae that mostly infect insects of the order Diptera (Litov et al., 2021; Longdon et al., 2010, 2012, 2015). Generally, sigmaviruses have a monopartite genome of around 12 kb to 15 kb (https://ictv.global/report/chapter/rhabdoviridae/rhabdoviridae/sigmavirus, accessed 19 February 2024) (Lefkowitz et al., 2018). In our study, the *Hermetia illucens* sigma-like virus 1 genome is split into two different transcripts (**Figure 1B**). In metatranscriptomic studies, it is not unusual to find fragments of viral genomes, which could be due to processing of the samples or errors introduced by the complexity of metatranscriptome assembly. All canonical sigmavirus genes were present across the two detected transcripts (**Figure 1B**), although we could not confirm that a large ORF on transcript 1 was the P gene based on sequence similarity, as it had no significant BLAST hits to other viruses or conserved domains. However, we propose that this is the P gene due to its position between the N gene and a partial M gene on this transcript (**Figure 1B**).

Sigmaviruses are used to study host-pathogen coevolution in the model organism *Drosophila melanogaster* (Duxbury et al., 2019; Fleuriet, 1996; Wayne et al., 2011; Yampolsky et al., 1999). *Drosophila* sigmaviruses are vertically transmitted by both males and females, and while *Drosophila* sigmaviruses are not known to be lethal to its host, they could be of interest to BSF breeders as they are associated with negative fitness effects such as delayed development and decreased fecundity (Fleuriet, 1981, 1996). However, not all effects of viral infection may be negative. Interestingly, one study found that sigmavirus infection in *Drosophila* positively influenced male reproductive success (Rittschof et al., 2013).

### 4.2 *Hermetia illucens* toti-like virus 2

Another novel virus detected in this study, which we call *Hermetia illucens* toti-like virus 2, groups within a large clade of insect-infecting toti-like viruses, and clusters closely with toti-like viruses that infect other insects of the order Diptera (Shi et al., 2016; W. Zhang et al., 2022). Totiviruses are a family of double-stranded RNA viruses that have historically been associated with fungi, but more recently, many metatranscriptomic studies are detecting toti-like viruses in other taxa, including insects (Giovannini et al., 2022; Huang et al., 2018; Koyama et al., 2015, 2016; Shi et al., 2016; Tighe et al., 2022; P. Zhang et al., 2018). *Hermetia illucens* toti-like virus 2 is the second toti-like virus discovered in BSF (Pienaar et al., 2022). In our phylogeny, *Hermetia illucens* toti-like virus 1 and 2 are separate from each other in distinct clades, where *Hermetia illucens* toti-like virus 1 groups with a virus that infects parasitoid wasps, and *Hermetia illucens* toti-like virus 2 groups with two viruses that infect other dipteran insects (**Figure 2B**).

### 4.3 Known BSF viruses

The two known viruses that we detected in this study occurred together very frequently, which is consistent with what we observed in our previous study (Walt et al., 2023). These samples were mostly detected in frass samples (**Figure 3**), which is also consistent with our previous study. Interestingly, a portion of BSF uncharacterized bunya-like virus genome was detected in a sample where no larvae were present, opening the question of whether BSF is the true host of this virus, or if it infects a component of their diet. Bunyaviruses can infect a wide variety of species, but our previous study showed that BSF uncharacterized bunya-like virus was most closely related to insect-infecting viruses. (Walt et al., 2023). Alternatively, this could be the result of cross contamination of samples.

### 4.4 Gene expression in putative antiviral genes in BSF

Although metatranscriptomics is useful for virus discovery and surveillance, it does not provide any information about the effects of viral infection on the host. To understand the effects that viruses may have on BSF, we analyzed the host transcriptome of BSF samples that were classified as either “virus-present” or “virus absent”. From a compiled table of 210 candidate genes (**Supplementary Table 1**) involved in BSF response to viruses, eight of them were differentially expressed. Of these, six are associated with the signaling pathways Imd and JAK/STAT (**Figure 4**), both of which can trigger antiviral immune responses (Costa et al., 2009; Dostert et al., 2005). One of the BSF genes that is significantly upregulated in virus-positive samples is orthologous to vir-1, a gene regulated by the JAK/Stat pathway that is also upregulated in *Drosophila* when infected by viruses (**Figure 4**, **Table 1**) (Carpenter et al., 2009; Dostert et al., 2005). Interestingly, we did not find significant differences in expression between orthologous components of the RNAi and PIWI-interacting RNA pathways (piRNA), both of which are important antiviral pathways in various dipteran species (Rosendo Machado et al., 2021). However, our sequencing datasets did not allow for the detection of small RNAs, the central molecules in the recognition of foreign nucleic acid. Evaluating the small RNA repertoire of BSF could provide more insight into the activation of antiviral pathways in this system (Ballinger et al., 2022).

Understanding the viruses routinely associated with BSF could be helpful in the event of a viral epidemic, which could cause detrimental impacts to the BSF industry. Our metatranscriptomic approach can detect known and novel viruses in BSF and frass and emphasizes the value of metatranscriptomic-based techniques to survey the viruses in BSF. Our approach also allowed the identification of host genes that could potentially be exploited as a screening tool of infection or fitness.

## Acknowledgments

The authors would like to thank the members of the Center for Environmental Sustainability through Insect Farming (CEIF) for their support in this work.

## Author Contributions

**H.K.W.** Conducted the analyses and wrote the original manuscript draft. **H.R.J., F.M.,** and **F.G.H.** acquired the funding, and provided supervision. All authors read and revised the final version of the manuscript.

## Data Availability

All new sequencing data generated in this study will be deposited in NCBI’s Sequence Read Archive (SRA) and will be available upon acceptance.

